# SuB3 a Simple Subcellular Fractionation Protocol for Localization Studies of Nucleic Acids and Proteins

**DOI:** 10.1101/2025.10.21.683662

**Authors:** Florencia S. Rodríguez, Lucas Servi, Leandro Quadrana, Ezequiel Petrillo

**Affiliations:** Instituto de Fisiología, Biología Molecular y Neurociencias (IFIBYNE), Buenos Aires, 1428, Argentina; Institute of Plant Sciences Paris-Saclay (IPS2), Université Paris-Saclay, CNRS, INRAe, Université Evry, Université Paris Diderot, 91190 Gif sur Yvette, France

## Abstract

Understanding the subcellular localization of RNA and proteins is critical to dissecting gene regulation in eukaryotic organisms. However, this task is elusive as existing fractionation methods often rely on protoplast isolation or commercial kits, that are labor-intensive, costly, and can introduce stress-induced transcriptomic and proteomic changes. Here, we present a simple, rapid, and cost-effective protocol for the fractionation of nuclear and cytoplasmic components directly from diverse plant tissues, that does not require protoplastization. This <u>Sub</u>cellular-fractionation protocol in <u>3</u> steps (a.k.a. “*bueno, bonito y barato”*- spanish for “good, nice and cheap”-), referred to as “SuB3”, yields nuclear- and cytoplasmic- enriched subcellular fractions suitable for downstream applications such as RT-PCR, RNA/cDNA sequencing, and Western blotting. The procedure is based on sequential detergent-assisted extraction and centrifugation and enables the simultaneous isolation of RNA and protein from the same biological material. Due to its simplicity, speed, and broad compatibility, this protocol is a valuable tool for plant molecular biology laboratories investigating subcellular dynamics of gene expression.

## Introduction

Subcellular localization of RNA and proteins is crucial for regulating gene expression and cellular function. Understanding where different molecules localize has key value to grasp their possible functions. Though this is relevant for general gene expression analyses, in the framework of alternative splicing regulation this is of utmost importance, as the role of a particular isoform, and its translability, can be easily inferred from its nuclear/cytoplasmic distribution. Intron retention is the most frequent alternative splicing event in plants (Marquez et al. 2012, 2015; Chaudhary et al. 2019; Petrillo 2023) and a possible destiny for transcripts that retain intron/s can be its nuclear retention (Göhring et al. 2014; Fuchs et al. 2021). Hence, to understand the biological consequences of alternative splicing regulation, knowing the subcellular localization of the generated isoforms is key (Tognacca et al. 2023). Similarly, protein localization is of key relevance to determine a protein’s role and to comprehend its turnover dynamics (Yousefi et al. 2021). Methods to rapidly and easily analyze nuclear/cytoplasmic localization of RNAs and proteins are then of great importance to understand the mechanisms and processes underlying cells and organism responses to environmental signals. Low costs, but more importantly, simplicity and speed (time saving/less RNA degradation), are relevant factors to be considered when analyzing if new methods are adoptable and easily reproducible in different labs around the globe.

In plants, analyzing the distribution of different factors in particular cellular compartments is very challenging, mainly due to the presence of a rigid cell wall. Traditional subcellular fractionation protocols often involve protoplast generation, which is not only time-consuming but can also introduce experimental artifacts due to stress responses (Barnes et al. 2019). To overcome these limitations, we have developed a simple protocol for rapid nuclear and cytoplasmic enrichment that eliminates the need for protoplastization and simplifies fractionation. Our protocol, termed **SuB3**, uses **liquid nitrogen–ground plant tissues, detergent-assisted cellular lysis**, and **3 sequential centrifugation steps** to obtain **RNAs** and/or **proteins** from **nuclear and cytoplasmic compartments**. In this work we describe the method in detail and briefly compare its performance against widely used protocols, demonstrating its effectiveness and simplicity.

## Material and Methods

### Plant Material

To evaluate the usefulness of the introduced protocol we used *Arabidopsis thaliana* (Col-0) two-week-old seedlings, *Solanum lycopersicum* one-week-old seedlings, and *Chenopodium quinoa* one-week-old seedlings.

For most experiments with *A. thaliana* seedlings, seeds were first stratified for three days in the dark at 4 °C and then germinated on Murashige and Skoog (MS) medium (pH 5.7, buffered with 2-(N-morpholino)ethanesulfonic acid (MES) supplemented with 1.5% agar. Seedlings were grown at 22 °C under constant white light with an irradiance of 70–100 μmol photons m^−2^ s^−1^.

Adult plants were grown in growth chambers at 22 °C or 25 °C depending on the species, under long-day conditions (16 h light/8 h dark). For these experiments, we used leaves from three-week-old *Arabidopsis thaliana* (Col-0) plants, three-week-old *Nicotiana tabacum* plants, and three-week-old *Solanum lycopersicum* plants.

### Lysis Buffer preparation

#### Stock solutions

1 M KCl. Make 50 mL. Filter through a 0.45 µM filter into a sterile bottle.

1 M MgCl_2_. Make 50 mL. Filter through a 0.45 µM filter into a sterile bottle.

1 M Tris-HCl (pH 7.5). Make 50 mL. Adjust pH with NaOH. Autoclave.

50% Glycerol. Autoclave.

10% NonidetNP40 (IGEPAL). Make 5mL with sterile water Deionized water. Autoclave.

#### Working solutions

### 1x Lysis Buffer

- 0.5 M sucrose (0.1712 g/mL weighed and added right before use)
- 50 mM KCl
- 5 mM MgCl_2_
- 50 mM Tris-HCl (pH 7.5)
- 25% Glycerol
- 0.1% IGEPAL (added right before use)

The 1x Lysis Buffer can be prepared up to 1 week in advance (without sucrose and IGEPAL) and stored for up to 1 month at −20°C. It is important to always keep it in the cold before use.

### Subcellular Fractionation with the suB3 Protocol

Starting material for both RNA and protein isolation should be over 250 mg of ground plant tissue. Specifically around 100 mg are used for “Total” RNA and/or protein isolation for proper comparisons, and the rest (∼150 mg) is used as starting material for the SuB3 Protocol:

***Step 1*)** resuspend ∼150 mg of ground tissue in 400 µL ice-cold Lysis Buffer. Mix thoroughly by inversion to homogenize and centrifuge at 3300 g for 3 minutes at 4 °C. **Pellet 1 should be considered as the source of the Nuclear-enriched Fraction. Supernatant 1 should be considered as the source of the Cytoplasmic-enriched Fraction**. NONE of them, supernatant or pellet from this first step of centrifugation, should be discarded! Following steps in the protocol are explained for each fraction separately (see Figure 1 for clarifications).

**Figure 1.**
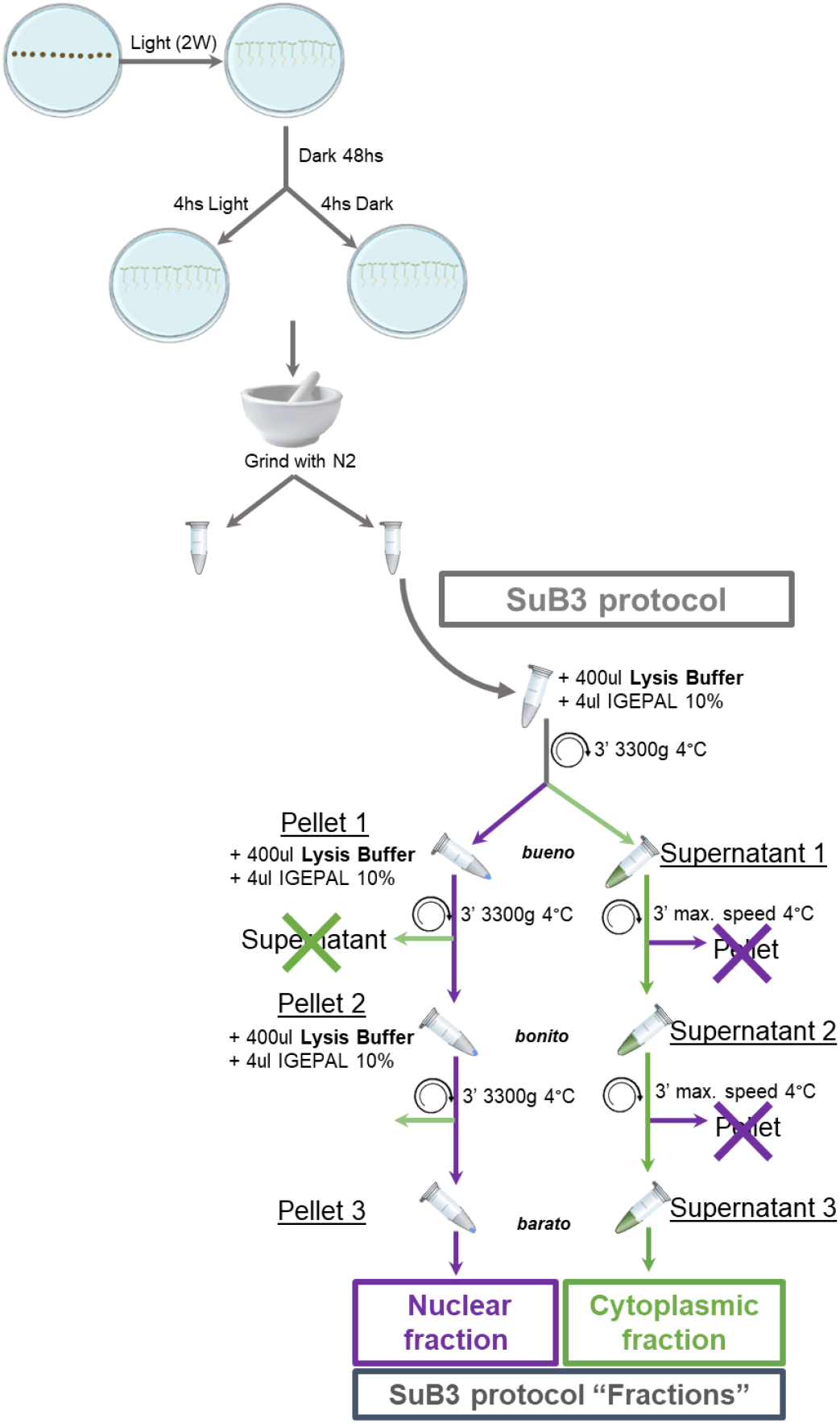
Schematic overview of the SuB3 protocol. Col-0 seeds were stratified for 2 days in darkness at 4 °C, germinated under continuous white light for 2 weeks (2W), and subsequently transferred to darkness for 48 h. At this stage, half of the plates were exposed to light while the remaining half were kept in darkness. These samples were used to establish the new fractionation protocol illustrated in the figure. The schematic shows the SuB3 workflow for obtaining total, nuclear, and cytoplasmic fractions from ground tissue.

***Nuclear-Enriched Fraction, from Pellet 1. Step 2*)** add 400 µL ice-cold Lysis Buffer to the tube containing Pellet 1. Carefully mix by inversion, avoiding air bubbles and keeping the samples as cold as possible. When the pellet cannot be resuspended by inversion, pipetting up and down is an option, however, to avoid nuclei disruption a wide-bore (or cutted) 1 mL pipette tip should be used. Centrifuge at 3300g for 3 minutes at 4°C.

***Step 3*)** carefully remove and discard supernatant without disturbing the Pellet 2. Add 400 µL ice-cold Lysis Buffer to the tube containing Pellet 2. Carefully mix by inversion, avoiding air bubbles and keeping the samples as cold as possible. Centrifuge at 3300g for 3 minutes at 4°C, then discard supernatant and **use Pellet 3 for further isolation of nucleic acids and/or proteins** (details in next section).

***Cytoplasmic-Enriched Fraction, from Supernatant 1. Step 2*)** centrifuge the supernatant from ***Step 1*** at 13000 g for 3 minutes at 4°C, transfer the Supernatant 2 to a new tube.

***Step 3*)** centrifuge the tube containing Supernatant 2 at 13000 g for 3 minutes at 4°C. Transfer the new supernatant, Supernatant 3, to a new tube. **Supernatant 3 is used for further isolation of nucleic acids and/or proteins** (details in next section).

### Chromatin and Nucleoplasm Isolation

In order to separate chromatin and nucleoplasm, SuB3 Pellet 3 was resuspended in 100 µL Nuclei Buffer. This suspension was processed following the protocol of Zhang *et al*. 2022 (*Chromatin extraction* section) until the Chromatin fraction was obtained as Pellet and Nucleoplasm fractions 1 and 2 as Supernatants.

### RNA and Protein Isolation

Proteins for Western Blot analyses can be directly obtained from the Total, Nuclear and Cytoplasmic fractions by adding protein extraction buffer directly to the proper “fractions”: ground tissue for Total; Pellet 3 for Nuclear and Supernatant 3 for Cytoplasmic. If proteins and RNAs are going to be evaluated, then we recommend using a reagent as Trizol (see below). When downstream analyses require purified RNAs, as for massive sequencing, then using a commercial kit as NucleoSpin RNA Plant (Macherey-Nagel), RNeasy Plant (Qiagen) or Spectrum Total RNA (Sigma) is recommended.

When analyzing isoform accumulation in different subcellular compartments by RT-PCR, Trizol (Sigma) extraction was performed following the manufacturer’s recommendations. Briefly, RNA can be isolated from “Total”, nuclear-enriched, and cytoplasmic-enriched samples by adding 1 mL TriPure/Trizol/TriReagent (or any other guanidinium-phenol based product) directly to the ground tissue (“Total”), to the nuclear pellet of the SuB3 protocol (Pellet 3), or to 200 µL of cytoplasmic-enriched fraction of the SuB3 protocol (Supernatant 3). In the latter case, if using an isolation reagent for Liquid Samples, volumes of supernatant can be increased, which would lead to higher yields for this fraction. After homogenization by vigorous shaking (rock the samples for 1 minute; avoid vortexing them as that may lead to fragmentation of nucleic acids), RNA can be extracted following manufacturers’ protocols without relevant modifications. While RNA is then precipitated from the aqueous phase after chloroform addition, proteins can be extracted from the organic phase via acetone precipitation. For protein precipitation, 350 µL of the organic phase are pipetted directly from the bottom of the tube avoiding the interphase. This volume is transferred to a 2 mL tube and 1.5 mL cold acetone (−20°C) are immediately added. Acetone must be used cold, stored at −20°C and used directly from the freezer. After homogenization by inversion, incubate the samples at −20°C overnight to improve the yield. After that period, following a centrifugation step, the pellet is washed once with ethanol 100 % and twice with ethanol 70 %. The pellet is then dried and resuspended 20 minutes at 60 °C directly in Laemmli’s buffer. For the experiments of the present manuscript, samples were sonicated 5 seconds at 15% amplitude using Sonic Dismembrator Model 500 (Fisher Scientific). The samples for Western blot analysis are ready after boiling for 5 minutes at 90°C.

For sequencing using Oxford Nanopore technologies RNAs are further purified using the NucleoSpin RNA Plant kit. For this, Total and Nuclear (Pellet 3) RNAs can be purified directly following the manufacturer’s instructions. For the Cytoplasmic fraction (Supernatant 3) the protocol was modified as follows, 350 µL of the Supernatant 3 (SuB3 Protocol) were directly loaded into the column of the kit in the step 2 “Lyse Cells” of their protocol without adding lysis buffer to the sample. RNA was processed following the ONT kit recommendations.

Note: RNA and protein extraction procedures should be decided based on downstream applications. Here we show some options, they work for RT-PCR-qPCR, Western Blots and 3rd generation sequencing, but it is up to the user to adapt the protocol and decide the best downstream purification protocol considering their particular questions or downstream methodologies.

## Results

### Determining the utility of the SuB3 Protocol

To assess the performance and versatility of our SuB3 protocol, we conducted a series of molecular analyses across RNA and protein fractions. These anlyses allowed us to benchmark subcellular enrichment and validate the method’s applicability across species and tissues.

When analyzing the RNA obtained from our Sub3-derived subcellular fractions, we observed that the relative amount of nuclear RNA is lower than that of cytoplasmic RNA, and both are lower than the Total fraction (Figure 2A). This is consistent with the multiple washing steps included in the protocol, which inevitably lead to material loss. Moreover, this is supporting specific fractionation, as ribosomal RNAs are known to accumulate preferentially in the cytoplasm, and are consistently highly depleted from the nuclear fraction. To validate our subcellular fractionation, we used two complementary approaches that allowed us benchmarking the protocol for RNAs and for proteins. First, we analyzed the alternative splicing pattern of the *At-SEF* transcripts by semi-quantitative (competitive) RT-PCR (Petrillo *et al*., 2014; Fuchs *et al*., 2021; Göhring *et al*., 2014). This gene has been previously characterized to produce three isoforms, two of which retain one or two introns and are predominantly nuclear (Göhring et al, 2014), and another one (fully spliced) that is rapidly exported to the cytoplasm and detected only to low levels in the nucleus. When analyzing the fractions from the SuB3 protocol, as expected, the fully spliced isoform is mainly detected in the cytoplasmic fraction, whereas the intron-retaining isoforms (*IR1 and IR1+2*) are enriched in the nuclear fraction (Figure 2A). Second, we analyzed specific proteins’ localization. Since proteins can also be purified from the organic phase of a TriPure reagent RNA extraction (see M&M section), we performed a western blot using antibodies against histone H3 as a nuclear marker and cytosolic fructose-1,6-bisphosphatase (cf-BFase) as a cytoplasmic marker. As shown in Figure 2B, the cytoplasmic fraction was indeed enriched in cf-BFase with barely undetectable histone levels, whereas the nuclear fraction displayed the opposite profile. In the Total fraction both proteins are detected, as expected. In addition, we performed Nanopore cDNA sequencing on the three fractions and specifically analyzed *At-SEF* gene to validate our observations using a 3rd generation sequencing independent approach (Figure 2C). Taken together, these results confirm that our protocol simply enables an efficient separation of nuclear and cytoplasmic fractions, and that both RNAs and proteins of suitable quality can be recovered from them for a wide range of downstream molecular analyses.

**Figure 2.**
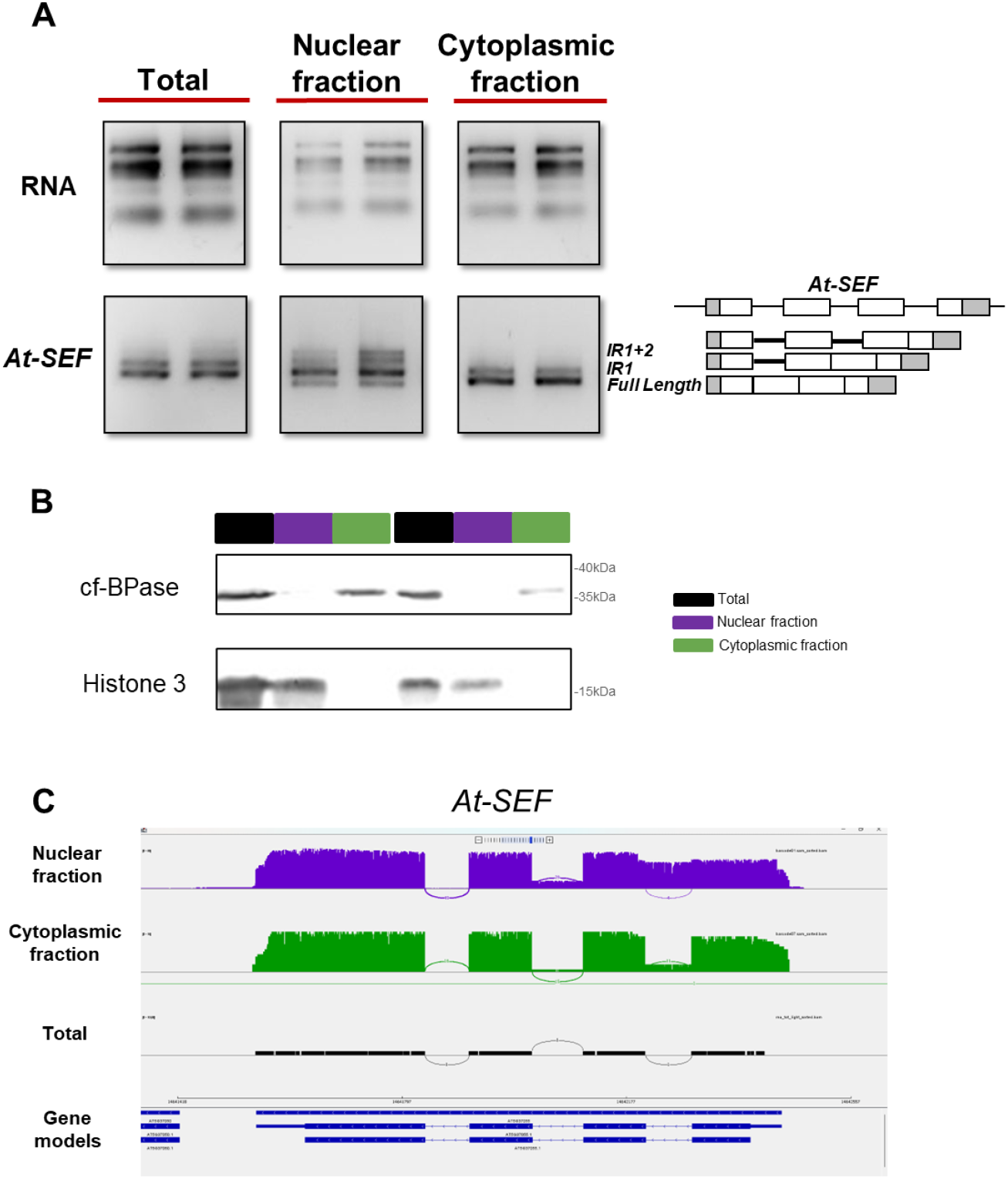
Validation of RNA and protein extración using the SuB3 protocol. (A) RNA extracted from SuB3 fractions. RNA integrity profiles for Total, Nuclear, and Cytoplasmic fractions. Validation of RNA purity was performed by RT–PCR using primers targeting *At-SEF* isoforms. Amplified products were resolved on 1.5% agarose gels. (B) Protein extraction from SuB3 fractions. Validation of protein enrichment in the same fractions by Western blot using fraction-specific antibodies: H3 (nuclear marker) and cf-BPase (cytoplasmic marker). (C) Validation by Nanopore cDNA sequencing across the three fractions. IGV visualization of *At-SEF* isoforms in Total, Nuclear, and Cytoplasmic fractions. The gene model structure is shown below.

To explore whether our nucleus-enriched fraction can be further processed to analyze chromatin associated and nucleoplasm-free transcripts, we implemented a modification of the SuB3 protocol (see M&M) and compared it with a published method to separate chromatin and nucleoplasmic fractions (Zhang *et al*. 2022). As shown in Figure 3, the *At-SEF* isoforms obtained from nuclei isolated with the Sub3 protocol can be further divided into RNAs associated to the nucleoplasmic fraction and those associated to chromatin, hence, the SuB3 Nuclear-enriched fraction is composed by whole nuclei. Moreover, the protocol can be easily modified to obtain further fractions from these whole nuclei (Figure 3).

**Figure 3.**
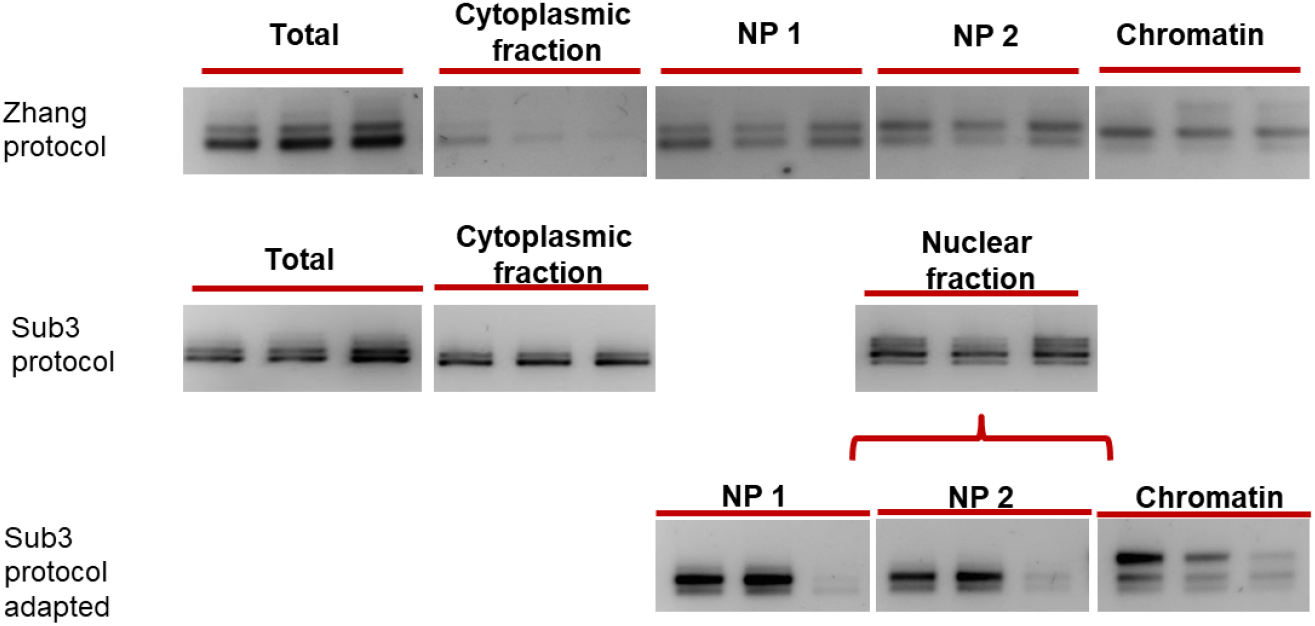
Modification of the SuB3 protocol for Chromatin and Nucleoplasmic separation. (A) Comparison between the chromatin isolation protocol described by Zhang et al. (2022) and the adapted SuB3 version. Validation of RNA separation was performed by RT–PCR using *At-SEF*-specific primers. Amplified products were resolved on 1.5% agarose gels.

Interestingly, using protoplasts to analyze subcellular fractions severely affects the alternative splicing patterns of *At-RS31* and *At-SR30*, among other splicing related factors, dramatically reducing the levels of the “longer” alternative isoforms (Petrillo *et al*., 2014; Fuchs *et al*., 2021). The nature of the SuB3 protocol, starting from plant tissues directly frozen in liquid nitrogen, avoids any perturbation of alternative splicing patterns due to protoplast preparation. To further assess this we evaluated the two mentioned genes, known to undergo alternative splicing and to generate isoforms with potential nuclear retention, by RT-PCR in the SuB3 generated fractions. As shown in Figure 4, intron-retention isoforms were preferentially enriched in the nucleus, whereas the coding isoforms were detected both in the nucleus and in the cytoplasm, with higher abundance in the cytoplasmic fraction as expected. Remarkably, all the fractions arising from the SuB3 protocol show higher levels of the longer alternative isoforms for these genes when compared to RNA isolated from protoplasts (Fig. 4A). In addition, we performed Nanopore sequencing on the three fractions and specifically searched for these genes to validate our observations using an independent approach. As shown in the IGV browser, the results were consistent with our RT–PCR analyses, confirming that the method is suitable not only for isoform distribution analysis by RT–PCR but also highly valuable for third generation sequencing-based approaches (Fig. 4B).

**Figure 4.**
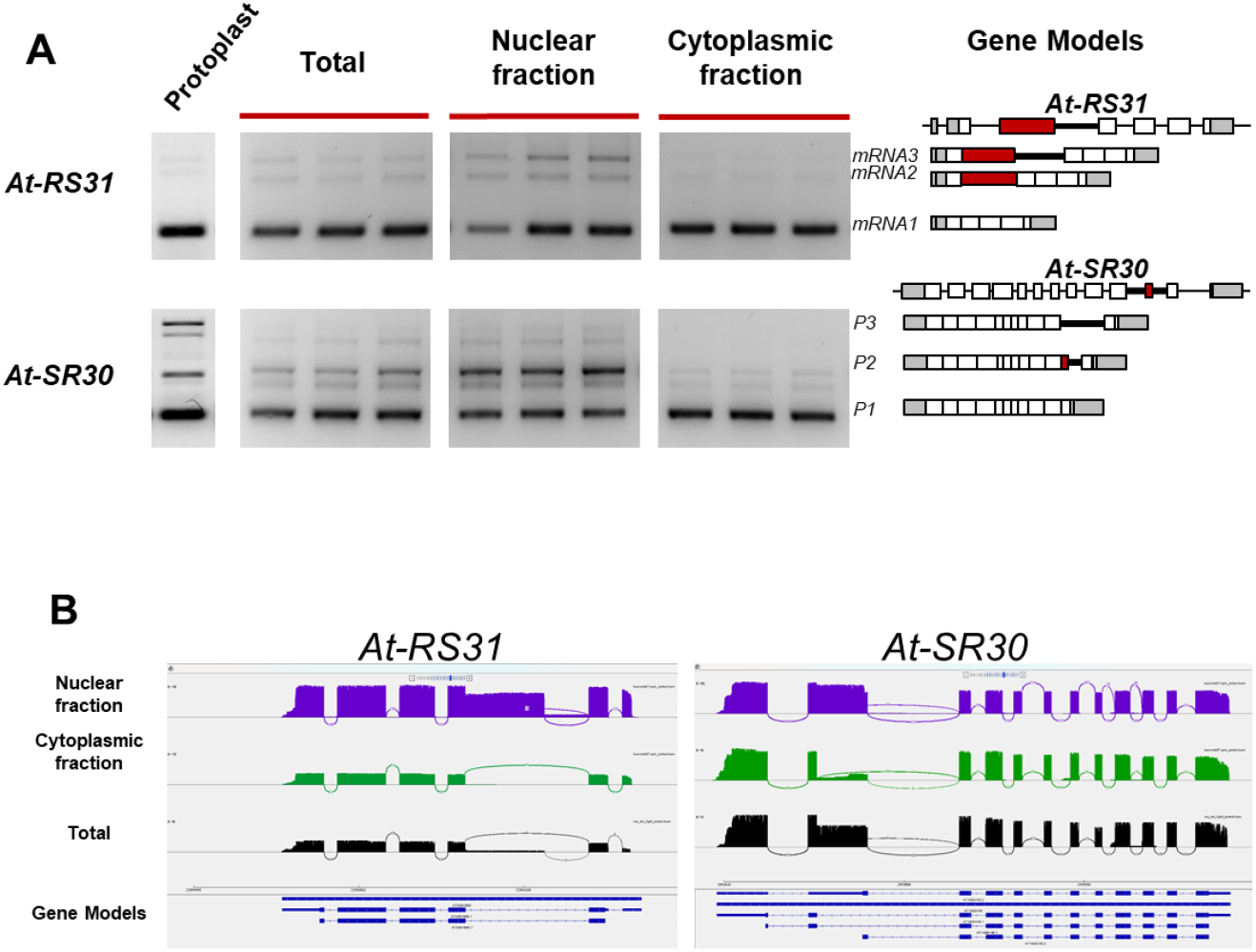
Subcellular localization of *At-RS31* and *At-SR30* isoforms. (A) Alternative splicing patterns from Protoplast, Total, Nuclear-, and Cytoplasmic fractions were analyzed by RT–PCR. Amplified products were resolved on 1.5% agarose gels. The right panel shows the corresponding gene models and isoforms for each gene. For Protoplast samples, different SR30 splicing primers were used to visualize the longer isoforms. (B) Nanopore cDNA sequencing of the three fractions, focusing on *At-RS31* and *At-SR30*. IGV visualizations show isoform distribution in Total, Nuclear, and Cytoplasmic fractions along with their gene model structures.

Finally, we tested whether our protocol could be used with other tissues and other plant species beyond *Arabidopsis thaliana*, including models widely used in molecular plant biology. Given that protein conservation is generally higher than RNA sequence conservation, we chose to validate the fractionation by protein extraction followed by western blotting with H3 (nuclear marker) and cf-BPase (cytoplasmic marker). We found that, starting either from adult leaves or seedlings and adjusting the amount of tissue powder accordingly, we successfully obtained ours three enriched fractions in at least four different species (*Arabidopsis thaliana, Nicotiana benthamiana, Solanum lycopersicum* and *Chenopodium quinoa*) (Figure 5).

**Figure 5.**
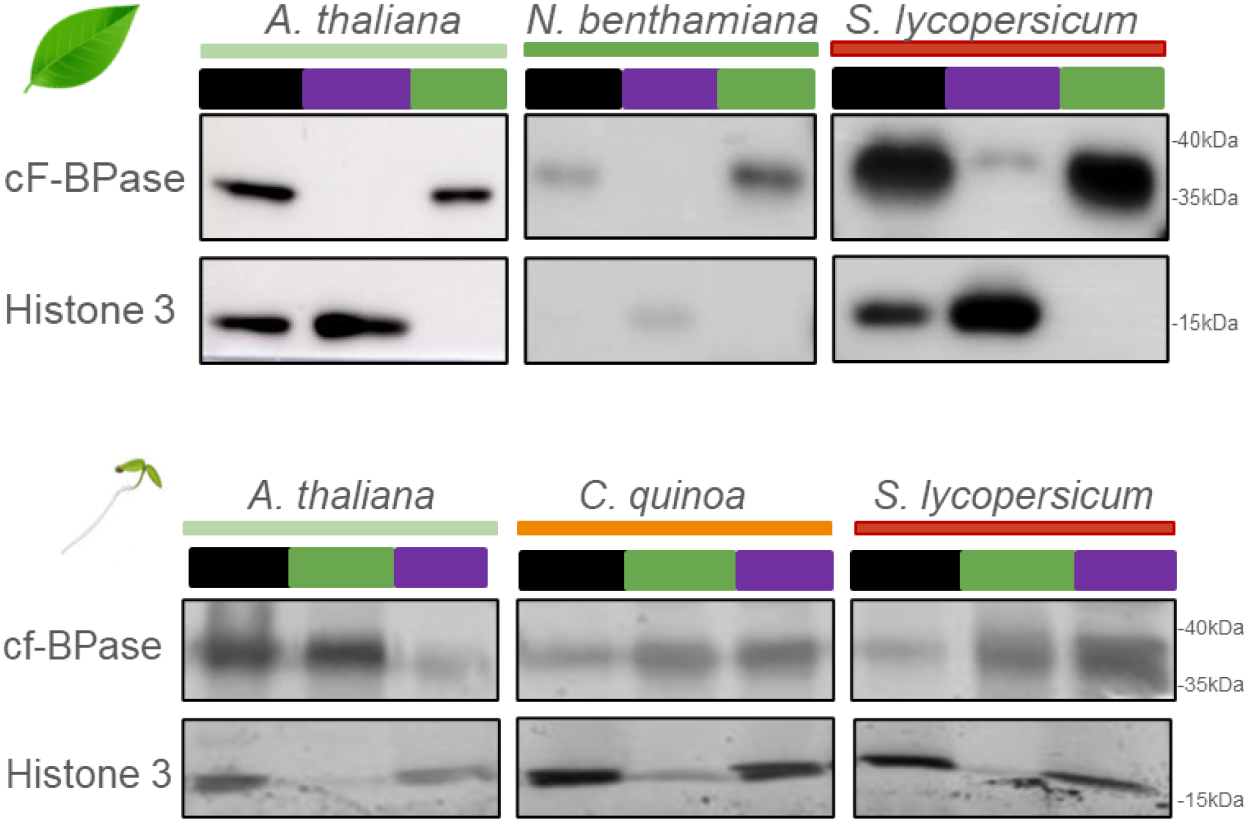
The SuB3 protocol is applicable to different species and tissues. (A) Protein extraction from SuB3 fractions using leaves of three adult plant species (*Arabidopsis thaliana, Nicotiana benthamiana*, and *Solanum lycopersicum*). Protein enrichment validation by Western blot using fraction-specific antibodies: H3 (nuclear marker) and cfBPhase (cytoplasmic marker). (B) Protein extraction from SuB3 fractions using seedlings of three plant species (*Arabidopsis thaliana, Chenopodium quinoa*, and *Solanum lycopersicum*). Validation of protein enrichment by Western blot using the same fraction-specific antibodies: H3 (nuclear marker) and cf-BPase (cytoplasmic marker).

These results strongly suggest that the protocol could be extended to many other species and potentially applied to more complex tissues such as seeds or fruits. This versatility highlights the broad applicability of the protocol as a robust and cost-effective tool to investigate subcellular RNA and protein localization across diverse plant systems.

## Discussion

The SuB3 method presents a reliable and accessible approach to nuclear and cytoplasmic fractionation (Table I). It eliminates the need and caveats of using protoplasts, minimizing artifacts and reducing cost and time. While not as pure as FACS- or INTACT-based approaches, the method’s balance of simplicity and effectiveness makes it well-suited for routine molecular biology applications in plant research. A key but not always addressed question in the alternative splicing field is what is the fate of the different isoforms, this method allows to simply know if an isoform of interest has the potential to be translated, easily showing nuclear retention or pointing to possible degradation in the cytoplasm. This is a fundamental issue when studying the outcomes of alternative splicing and the functions of different transcripts aiming to understand their impact on an organism’s phenotype (Tognacca *et al*., 2023).Furthermore, the SuB3 protocol can be extended to interrogate about the precise nuclear accumulation of the transcripts, and know if the isoforms are chromatin-bound or present in the nucleoplasm.

**Table 1.**
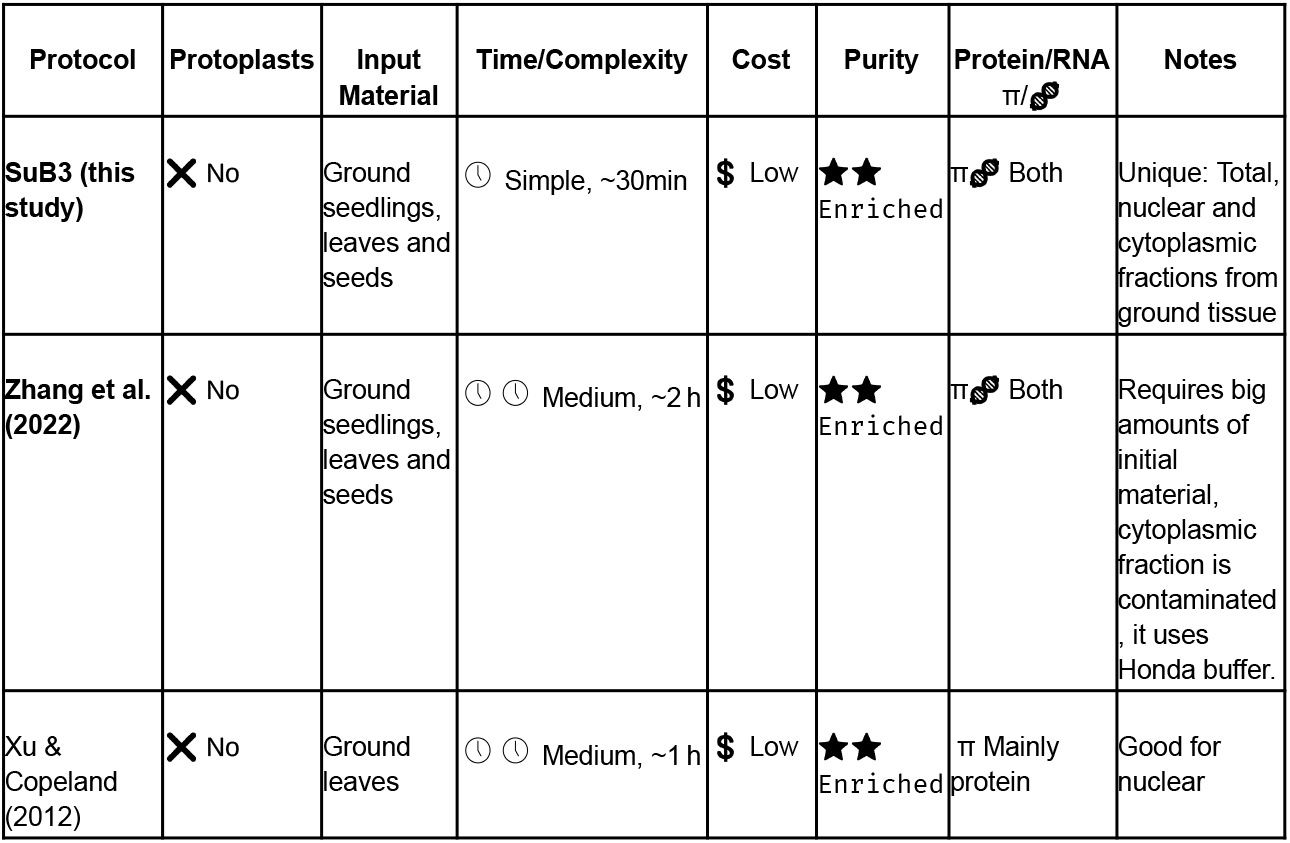
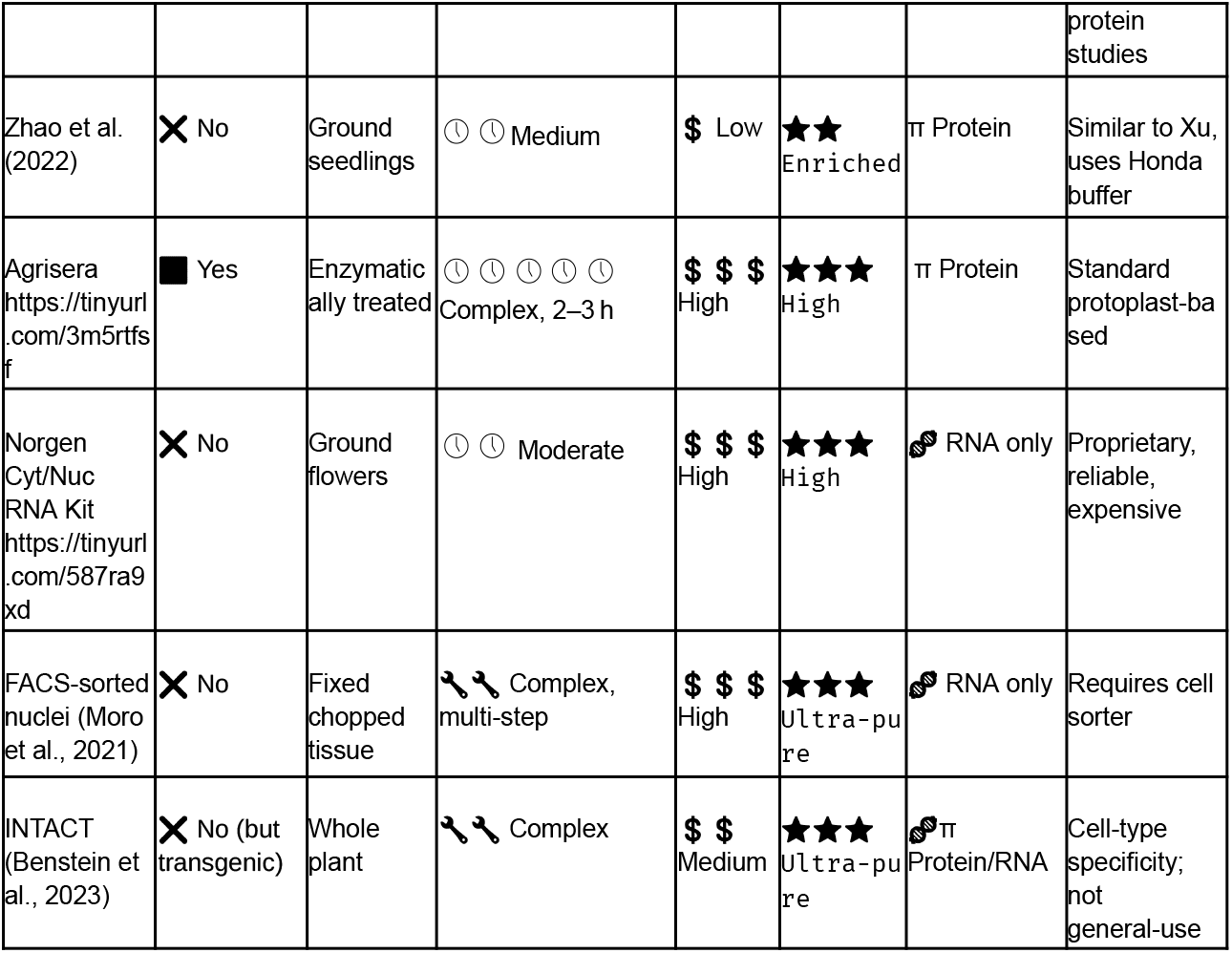
Comparison of the SuB3 protocol with previously published methods for nuclear and cytoplasmic fractionation. Summary of key features of the SuB3 protocol in comparison with other available methods for plant subcellular fractionation. The table highlights differences in required material, time, cost, and the possibility of recovering both RNA and proteins from the same sample.

**Table 2.**
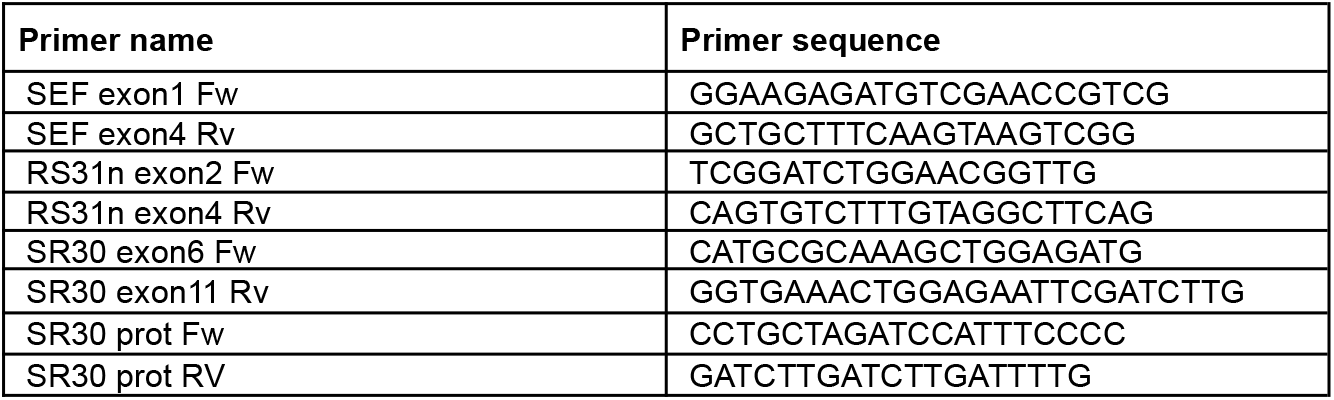
Primers used in the manuscript.

## Acknowledgments

We thank the whole community at IFIBYNE and IPS2 for the exciting and inspiring working atmosphere. We do not thank the lack of support from argentinian authorities, the lack of vision and the sad degradation of the scientific system in our country. We specially thank the ICGEB as their CRP/ARG22-03 grant (to EP) is the only funding we have in our lab in Argentina, thanks for your support and for keeping our lab running in these dark times. As Bernardo Houssay once said, ‘Science is not expensive; expensive is ignorance.’ We echo this sentiment and advocate for sustained investment in research as a cornerstone of societal progress.

## Author Contributions

Ezequiel Petrillo conceived the manuscript and, together with Florencia S. Rodríguez, prepared and wrote the manuscript with input from Lucas Servi and Leandro Quadrana. FSR together with LS performed the experiments and generated the figures. All authors read and approved the final version of the manuscript.

## Data Availability

RNA-seq data shown in the manuscript are available from the lab and the whole dataset will be soon released in GEO.

